# Dynamic virulence-related regions of the fungal plant pathogen *Verticillium dahliae* display remarkably enhanced sequence conservation

**DOI:** 10.1101/277558

**Authors:** Jasper R.L. Depotter, Xiaoqian Shi-Kunne, Hélène Missonnier, Tingli Liu, Luigi Faino, Grardy C.M. van den Berg, Thomas A. Wood, Baolong Zhang, Alban Jacques, Michael F. Seidl, Bart P.H.J. Thomma

## Abstract

Selection pressure impacts genomes unevenly, as different genes adapt with differential speed to establish an organism’s optimal fitness. Plant pathogens co-evolve with their hosts, which implies continuously adaption to evade host immunity. Effectors are secreted proteins that mediate immunity evasion, but may also typically become recognized by host immune receptors. To facilitate effector repertoire alterations, in many pathogens, effector genes reside in dynamic genomic regions that are thought to display accelerated evolution, a phenomenon that is captured by the two-speed genome hypothesis. The genome of the vascular wilt pathogen *Verticillium dahliae* has been proposed to obey to a similar two-speed regime with dynamic, lineage-specific regions that are characterized by genomic rearrangements, increased transposable element activity and enrichment in *in planta*-induced effector genes. However, little is known of the origin of, and sequence diversification within, these lineage-specific regions. Based on comparative genomics among *Verticillium* spp. we now show differential sequence divergence between core and lineage-specific genomic regions of *V. dahliae*. Surprisingly, we observed that lineage-specific regions display markedly increased sequence conservation. Since single nucleotide diversity is reduced in these regions, host adaptation seems to be merely achieved through presence/absence polymorphisms. Increased sequence conservation of genomic regions important for pathogenicity is an unprecedented finding for filamentous plant pathogens and signifies the diversity of genomic dynamics in host-pathogen co-evolution.

## INTRODUCTION

Numerous microbes engage in symbiotic relationships with plants, comprising beneficial, commensalistic and parasitic relationships where each partner evolves towards its optimal fitness. Consequently, parasitic interactions between plants and microbial pathogens evolve as arms races in which plants try to halt microbial ingress while pathogens strive for continued symbiosis (Jones and Dangl 2006; Thomma et al. 2011; Cook et al. 2015). In such arms races, plant pathogens evolve repertoires of effector proteins, many of which deregulate host immunity, to enable host colonization (de Jonge et al. 2011; Rovenich et al. 2014). Plants, in turn, evolve immune receptors that recognize various molecular patterns that betray microbial invasion; so-called invasion patterns that can also include effectors (Cook et al. 2015). Consequently, pathogen effector repertoires are typically subject to selective forces that often result in rapid diversification.

Effector genes are often not randomly organized in genomes of filamentous plant pathogens (Dong et al. 2015). For instance, effector genes of the potato late blight pathogen *Phytophthora infestans* reside in repeat-rich regions that display increased structural polymorphisms and enhanced levels of positive selection (Haas et al. 2009; Raffaele et al. 2010). Based on this and observations in other pathogenic species, it has been proposed that many pathogens have a bipartite genome architecture with essential household genes residing in the core genome and effector genes co-localizing in repeat-rich compartments; a phenomenon that has been coined a two-speed genome (Croll and McDonald 2012; Raffaele and Kamoun 2012; Seidl and Thomma 2017). Conceivably, such genome compartmentalization increases the evolutionary efficiency as basal functions of core genes are “shielded off” from increased evolutionary dynamics that rapidly diversify effector gene repertoires. Repeat-rich genome regions display signs of such accelerated evolution as they are often enriched for structural variations such as presence/absence polymorphisms (Raffaele et al. 2010) or chromosomal rearrangements (de Jonge et al. 2013; Faino et al. 2016). In addition, increased diversification is also displayed on sequence levels in the form of higher substitution rates (Cuomo et al. 2007; van de Wouw et al. 2010) with a higher fraction of non-synonymous substitutions in genes located in repeat-rich regions compared to core genes (Raffaele et al. 2010; Stukenbrock et al. 2010; Sperschneider et al. 2015).

*Verticillium* is a genus of Ascomycete fungi, containing notorious plant pathogens of numerous crops, including tomato, cotton, olive and oilseed rape (Inderbitzin and Subbarao 2014). *Verticillium* spp. are soil-borne fungi that infect their hosts via the roots and then colonize xylem vessels, resulting in vascular occlusion by host depositions and by the physical presence of the pathogen itself (Fradin and Thomma 2006). Currently, ten *Verticillium* species are described (Inderbitzin et al. 2011a). All these *Verticillium* spp. are haploids, except for *V. longisporum* that is an interspecific hybrid that contains approximately twice the amount of genetic material of haploid *Verticillium* spp. (Inderbitzin et al. 2011b; Depotter et al. 2017). *V. dahliae* is the most notorious plant pathogen within the *Verticillium* genus, causing disease on hundreds of plant species (Inderbitzin and Subbarao 2014). Similarly, *V. albo-atrum, V. alfalfa, V. nonalfalfae* and *V. longisporum* are pathogenic, albeit with more confined host ranges (Inderbitzin et al. 2011a; Inderbitzin and Subbarao 2014). The remaining *Verticillium* spp., namely *V. isaacii, V. klebahnii, V. nubilum, V. tricorpus* and *V. zaregamsianum*, sporadically cause disease on plants and are considered opportunists with a mainly saprophytic life style rather than genuine plant pathogens (Ebihara et al. 2003; Inderbitzin et al. 2011a; Gurung et al. 2015).

*Verticillium* spp. are thought to have a predominant, if not exclusive, asexual reproduction as a sexual cycle has never been described for any of the species (Short et al. 2014). However, mating types, meiosis-specific genes and genomic recombination between clonal lineages have been observed for *V. dahliae*, suggesting that sexual reproduction is either cryptic or ancestral (Milgroom et al. 2014; Short et al. 2014). Nevertheless, mechanisms different from meiotic recombination contribute to the genomic diversity of *V. dahliae*, such as large-scale genomic rearrangements, horizontal gene transfer and transposable element (TE) activity (de Jonge et al. 2012; de Jonge et al. 2013; Seidl and Thomma 2014; Faino et al. 2016). Signs of these evolutionary mechanisms converge on particular genomic regions that are enriched in repeats and lineage-specific (LS) sequences (Klosterman et al. 2011; de Jonge et al. 2013; Faino et al. 2016). Intriguingly, also *in planta*-induced effector genes are enriched in these LS regions (de Jonge et al. 2013).

We previously reported that LS regions of *V. dahliae* are largely derived from segmental duplications (Faino et al. 2016). Gene duplications are important sources for functional diversification (Magadum et al. 2013), and thus here we aim to investigate whether and how the nucleotide sequences within the LS regions diverge. To this end, comparative genomics was performed across the *Verticillium* genus, to identify genomic regions showing accelerated and reduced rates of sequence diversification to further characterize the two-speed genome of *V. dahliae*.

## RESULTS

### LS sequences reside in four regions of the genome of *V. dahliae* strain JR2

Previously, we identified regions in the genome sequence of *V. dahliae* isolate JR2 that lack synteny to various other *V. dahliae* strains, including the completely sequenced genome of strain VdLs17, and thus these regions have been referred to as LS regions (Faino et al. 2015; Faino et al. 2016). In *V. dahliae* isolate JR2, the majority of these LS sequences cluster into four genomic regions; chromosomes 2 and 4 each contain one LS region, while two distinct LS regions reside on chromosome 5. To further characterize these LS regions, we here pursued high-quality genome assemblies of additional *V. dahliae* strains based on single-molecule real-time (SMRT) using the PacBio RSII system. Since *V. dahliae* strains JR2 and VdLs17 only recently diverged (de Jonge et al. 2013), we selected two *V. dahliae* strains that are more diverged (Supplemental_Figure_S1), namely strains CQ2 and 85S isolated from cotton in China and sunflower in France, respectively. We generated 430,378 (~110x coverage) and 500,428 (~130x coverage) filtered sub-reads for strains CQ2 and 85S, respectively, that were assembled into 17 and 40 contigs with a total size of 35.8 and 35.9 Mb, respectively (Supplemental_Table_S1), which is similar to the telomere-to-telomere assemblies of strains JR2 (36.2 Mb) and VdLs17 (36.0 Mb) (Faino et al. 2015). Subsequently, the assemblies of strains VdLs17, CQ2 and 85S were individually aligned to JR2 assembly. In total, 2.0%, 7.1% and 6.6% of the JR2 genome was not covered by sequences from VdLs17, CQ2 and 85S respectively, and 1.4% of the JR2 genome sequence could not be identified in any of the three other *V. dahliae* strains. The vast majority (88%, 82% and 91% for VdLs17, CQ2 and 85S, respectively) of these JR2 sequences without alignment is located in the previously identified four LS regions (Figure 1). Thus, despite the addition of more diverged *V. dahliae* strains, intraspecific presence/absence polymorphisms converge on the four previously identified genomic regions that are thus significantly more dynamic than other parts of the genome.

**Figure 1:**
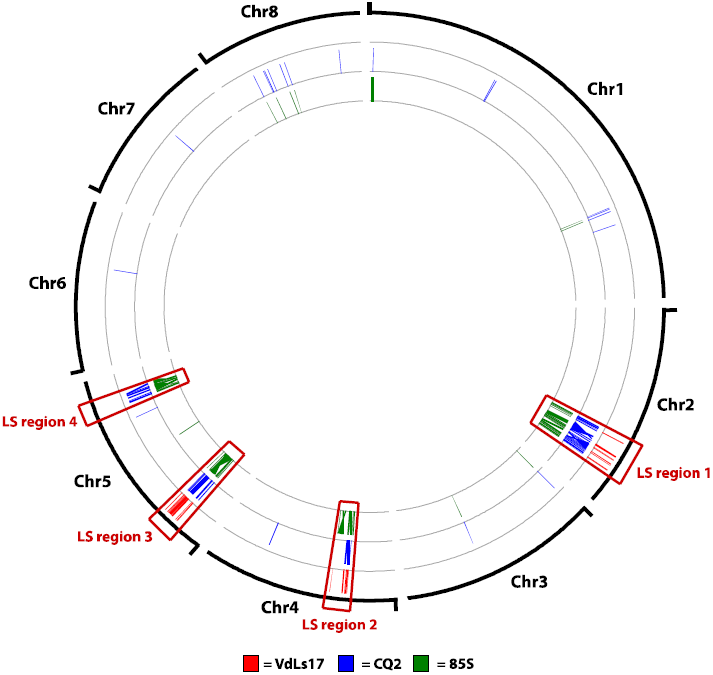
Locations of lineage-specific (LS) regions in the genome of *V. dahliae* strain JR2. LS regions were determined by individual comparisons to *V. dahliae* strains VdLs17 (red), CQ2 (blue) and 85S (green). Sequences of minimum 7.5 kb without an alignment to at least one of the other isolates are depicted at their respective position on the *V. dahliae* strain JR2 genome.

### LS regions share increased sequence identity to other *Verticillium* spp

To study interspecific sequence conservation, we aligned sequences of the phylogenetically closely related and previously sequenced *V. nonalfalfae* strain TAB2 (Jelen et al. 2016; Shi-Kunne et al. in preparation) to the genome assembly of *V. dahliae* strain JR2. While most of the genome of *V. dahliae* JR2 aligns with *V. nonalfalfae* strain TAB2 with a genome-wide average sequence identity of ~92%, particular genomic regions display an increased sequence identity, even up to 100% (Supplemental_Figure_S2). Intriguingly, these regions co-localize with the LS regions of *V. dahliae* strain JR2, implying that these LS regions are either derived from a recent horizontal transfer, subject to negative selection that depletes sequence polymorphisms, or encounter lower mutation rates.

In order to evaluate whether LS sequences also display increased sequence conservation when compared with other *Verticillium* spp., we aligned sequences of all other haploid *Verticillium* spp. to the *V. dahliae* JR2 genome assembly. These genomic data were previously generated (Jelen et al. 2016; Shi-Kunne et al. in preparation), apart from the data for *V. isaacii* strain PD660 that we sequenced using the Illumina HiSeq2000 platform (Supplemental_Table_S2). Genomic sequences of *V. dahliae* (windows of 500 bp) were aligned to the other *Verticillium* spp., displaying median identities ranging from 88 to 95% (Table 1). These percentages correspond to the phylogenetic distance of the respective species to *V. dahliae*. Sequence identities were similarly calculated in windows for the LS regions. Intriguingly, the LS regions displayed significantly increased sequence identities when compared with the core genome (Figure 2, Table 1), ranging from 92.3% median sequence identity for *V. zaregamsianum*, which is one of the phylogenetically most distantly related species to *V. dahliae*, to 100% median sequence identity for *V. alfalfae* and *V. nonalfalfae*. Thus, based on the genus-wide occurrence and a differential degree of sequence identity that reflects the phylogenetic distance to *V. dahliae*, we conclude that LS regions display increased sequence conservation when compared with the core genome, rather than originate form horizontal transfer events.

**Table 1:** **Sequence identities between *V. dahliae* (JR2) and other haploid *Verticillium* species (excluding repetitive regions).**

| Species/strain | Genome-wide<sup>a</sup><br>(%) | LS regions<sup>a</sup><br>(%) | # windows aligned to<br>LS regions<sup>b</sup> | *p*-value<sup>c</sup> |
| --- | --- | --- | --- | --- |
| *V. albo-atrum*/PD747 | 88.8 (0.12) | 97.6 (0.31) | 146 | 2.2e-16 |
| *V. alfalfae*/PD683 | 94.6 (0.03) | 100.0 (0.16) | 465 | 2.2e-16 |
| *V. nonalfalfae*/TAB2 | 94.8 (0.03) | 100.0 (0.09) | 1037 | 2.2e-16 |
| *V. nubilum*/PD621 | 88.4 (0.09) | 95.6 (0.41) | 144 | 2.2e-16 |
| *V. tricorpus*/PD593 | 89.0 (0.12) | 97.2 (0.32) | 162 | 2.2e-16 |
| *V. isaacii*/PD660 | 88.8 (0.12) | 98.6 (0.32) | 189 | 2.2e-16 |
| *V. klebahnii*/PD401 | 88.8 (0.11) | 97.8 (0.57) | 87 | 2.2e-16 |
| *V. zaregamsianum*/PD739 | 88.8 (0.10) | 92.3 (0.69) | 38 | 1.56e-5 |

<sup>a</sup>The percentage is the median sequence identity and the number between the brackets is the standard error.
<sup>b</sup>Windows of 500bp
<sup>c</sup>The *p*-value was calculated with a two-sided Wilcoxon rank-sum test

**Figure 2:**
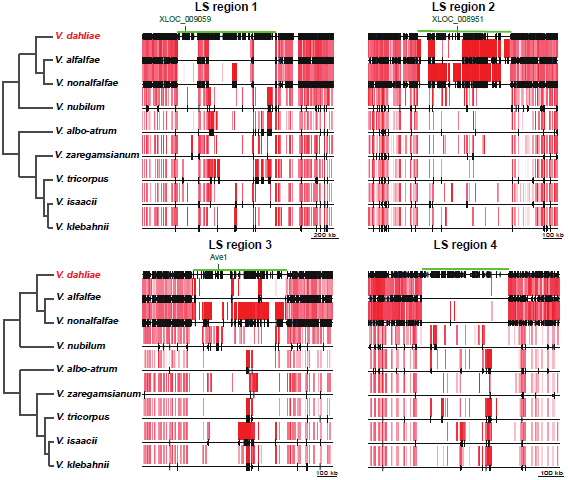
Interspecific alignments and sequence identity within and immediately adjacent to lineage-specific (LS) regions of *V. dahliae*. The green line indicates the extent of the LS region. The pink/red bars are *Verticillium* sequences alignments to JR2, whereas the intensity of their colour represents relative sequence identity for every *Verticillium* spp. individually (higher identity = red, lower identity = pink). The black, vertical stripes on the synteny lines represent predicted gene positions. For *V. dahliae* JR2, all predicted genes are depicted, whereas for other species only genes are depicted if these were successfully aligned. Locations of characterized *V. dahliae* effector genes are indicated: *Ave1, XLOC_008951* and *XLOC_009059* (de Jonge et al. 2012; de Jonge et al. 2013). Strains used in this figure: *V. dahliae* JR2, *V. alfalfae* PD683, *V. nonalfalfae* TAB2, *V. nubilum* PD621, *V. albo-atrum* PD747, *V. zaregamsianum* PD739, *V. tricorpus* PD593, *V. isaacii* PD660 and *V. klebahnii* PD401.

In order to evaluate whether the increased sequence conservation is specific only to LS regions, we aligned *Verticillium* sequences of high identity to the complete *V. dahliae* JR2 genome. For several species we used multiple strains at this stage. While some of these strains were previously sequenced (Supplemental_Table_S2) (Seidl et al. 2015; Jelen et al. 2016; Shi-Kunne et al. in preparation) others were newly sequenced (*V. albo-atrum* strain PD670, *V. klebahnii* strain PD659 and *V. zaregamsianum* strain PD736) using the Illumina HiSeq2000 platform. Nearly all (99-100%) of the *V. alfalfae* and *V. nonalfalfae* sequences that display >96% identity to *V. dahliae* strain JR2 sequenced localized in LS regions (Figure 3). Similarly, sequences of at least 100 kb with >90% identity of other *Verticillium* spp. mapped to *V. dahliae* JR2 LS regions, ranging from 70% in *V. nubilum* PD621 tot 95% in *V. albo-atrum* PD670 and *V. tricorpus* PD593 (Figure 3, Supplemental_Table_S3). In conclusion, increased sequence conservation is a genomic feature that is specifically associated with LS regions in *V. dahliae*.

**Figure 3:**
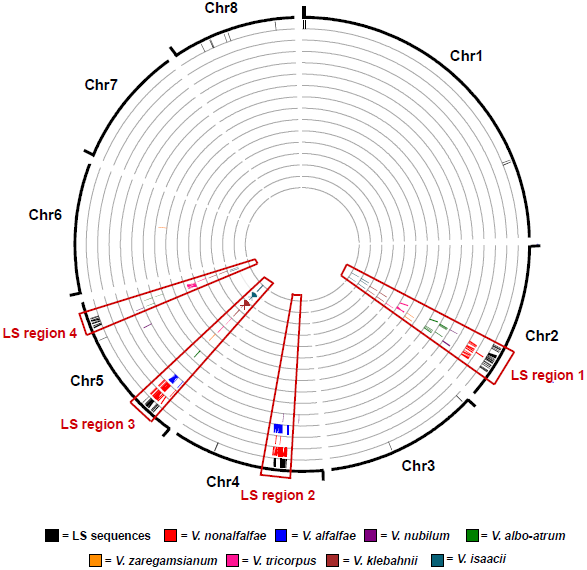
Regions of particular high sequence identity between *V. dahliae* and other haploid *Verticillium* species. Black bars correspond to lineage-specific sequences of *V. dahliae* strain JR2 (for details, see Figure 1). Sequences (≥7.5 kb) with high sequence identity in any of the other *Verticillium* spp. (≥96% for *V. alfalfae* and *V. nonalfalfae*, ≥90% for all other *Verticillium* spp.) are plotted at the corresponding position on the genome of *V. dahliae* strain JR2. Plotting order for the different *Verticillium* strains from the outside to inside of the circle: *V. dahliae* JR2 *V. nonalfalfae* TAB2 and Rec, *V. alfalfae* PD683, *V. nubilum* PD621, *V. albo-atrum* PD670 and PD747, *V. zaregamsianum* PD736 and PD739, *V. tricorpus* PD593 and MUCL9792, *V. klebahnii* PD401 and PD659, *V. isaacii* PD618 and PD660.

### LS regions with increased sequence conservation are not unique to *V. dahliae*

To investigate whether other *Verticillium* spp. similarly carry LS regions that display increased sequence conservation, we performed alignments using *V. tricorpus* strain PD593 as a reference because of its high degree of completeness, as seven of the nine scaffolds likely represent complete chromosomes (Supplemental_Table_S2) (Shi-Kunne et al. in preparation). Furthermore, this species belongs to the Flavexudans clade in contrast to *V. dahliae* that belongs to the Flavnonexudans clade. LS sequences of *V. tricorpus* strain PD593 were determined by comparison to *V. tricorpus* strain MUCL9792 (Seidl et al. 2015). In total, 98% of the PD593 genome could be aligned to MUCL9792. However, 48% of the sequences that are specific for *V. tricorpus* strain PD593 resided in one genomic region of 41 kb on scaffold 1 (Figure 4). Like for *V. dahliae* strain JR2, we were able to align sequences of other *Verticillium* spp. with high identity to the *V. tricorpus* strain PD593 genome (Figure 4): *V. isaacii, V. klebahnii* and *V. zaregamsianum* display a median genome identity of ~95% to *V. tricorpus*, while other haploid *Verticillium* spp. display ~88-89% median genome identity. Notably, regions that display significant higher sequence identify localized at the LS region on scaffold region, but also to an additional region of 23 kb on scaffold 6 (Figure 4, Supplemental_Figure_S3). For *Verticillium* strains with total alignments of at least 100 kb of high-identity sequences, the fraction of high-identity sequences that aligned to the scaffold 1 and 6 genome loci ranged from 49% for *V. nubilum* (PD621) up to 84% for *V. albo-atrum* (PD747) (Supplemental_Table_S4). As expected, the sequence identity to six of the eight other haploid *Verticillium* spp. was significantly higher in these two genome loci compared to the genome-wide median (Supplemental_Table_S5). No increase in sequence identity was found in alignments with *V. alfalfae* strain PD683 and *V. zaregamsianum* strain PD739 as only few regions with high sequence identity aligned to strain PD593 (Supplemental_Table_S4). Strains PD683 and PD739 only aligned 2 and 37 windows of 500 bp, respectively, to scaffold 1 and 6 loci of PD593 (Supplemental_Figure_S3, Supplemental_Table_S5). In conclusion, LS regions with increased sequence conservation are not unique to *V. dahliae*, but also occur in *V. tricorpus*, and thus likely in other *Verticillium* spp. as well.

**Figure 4:**
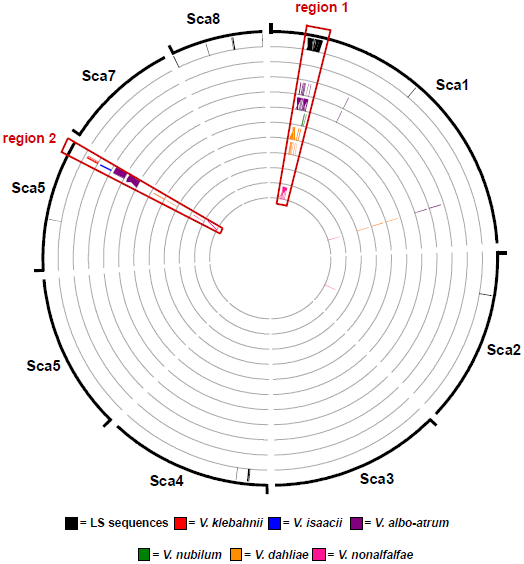
Regions of particular high sequence identity between *V. tricorpus* and other haploid *Verticillium* species. The eight biggest scaffolds of PD593 are depicted as these comprise over 99.5% of the genome. Black bars correspond to LS sequences (≥7.5 kb) in the PD593 genome without alignments to *V. tricorpus* strain MUCL9792. Sequences (≥7.5 kb) with relatively high sequence identity in any of the other *Verticillium* spp. (≥96% *V. isaacii, V. klebahnii* and *V. zaregamsianum*, ≥90% for all other *Verticillium* spp.) are plotted at the corresponding position on the genome of *V. tricorpus* strain PD593. Plotting order for the different *Verticillium* strains from the outside to inside of the circle: *V. tricorpus* PD593, *V. klebahnii* PD659, *V. isaacii* PD660, *V. albo-atrum* PD670 and PD747, *V. nubilum* PD621, *V. dahliae* JR2, VdLs17 and 85S, *V. nonalfalfae* TAB2 and Rec. Non-depicted *Verticillium* strains did not have sequences (≥7.5 kb) with previously mention identity to PD593.

### The pan-LS-genome distribution across the *Verticillium* genus

The increased conservation of LS sequences in two diverged *Verticillium* species indicates that the origin of many of the LS regions is ancestral and predates their speciation. Hence, we constructed a pan-LS-genome to determine the distribution of conserved sequences across the *Verticillium* genus. To compose a pan-LS-genome, we combined regions with increased sequence conservation of four *Verticillium* spp., namely *V. dahliae* strain JR2, *V. alfalfae* strain PD683, *V. tricorpus* strain PD593 and *V. klebahnii* strain PD401, motivated by their high assembly contiguity and distribution throughout the *Verticillium* genus (Inderbitzin et al. 2011a; Shi-Kunne et al. in preparation). After removal of repetitive and duplicated sequences, we obtained a pan-LS-genome of ~2 Mb, of which 60% occurs in genomes of clade Flavexudans and 72% in clade Flavnonexudans (clade pan-LS-genomes) (Figure 5). Next, the distribution of the pan-LS-genome and the clade pan-LS-genomes was evaluated for all *Verticillium* strains individually (Figure 5, Supplemental_Table_S6). The proportion of the LS-pan-genome differed markedly between *Verticillium* strains and ranged from 12% for *V. nubilum* strain PD621 up to 58% for *V. dahliae* strain JR2 (Figure 5). Notably, by using a limited number of isolates in the consensus reconstruction, retentions are likely biased towards strains that are phylogenetically closer related to the species that were used to compose the pan-genome: *V. alfalfa, V. dahliae, V. klebahnii* and *V. nonalfalfae*. However, *V. albo-atrum* strains contained considerably more of the pan-LS-genome compared to *V. zaregamsianum* and *V. isaacii* strains, despite its phylogenetically more distant relation to *V. klebahnii* and *V. tricorpus* (Figure 5). Moreover, LS contents do not only differ considerably between species but also within species as we also observed large intra-specific differences. For example, the genome of *V. dahliae* strain VdLs17 contains less than two thirds of the content present in the LS regions of the strain JR2 genome despite the recent divergence of the two strains (Figure 5, Supplemental_Figure_S1) (Faino et al. 2015). Thus, sequences with increased conservation are genus-wide associated with dynamic genomic regions of *Verticillium* spp. as their contents vary greatly between and within species.

**Figure 5:**
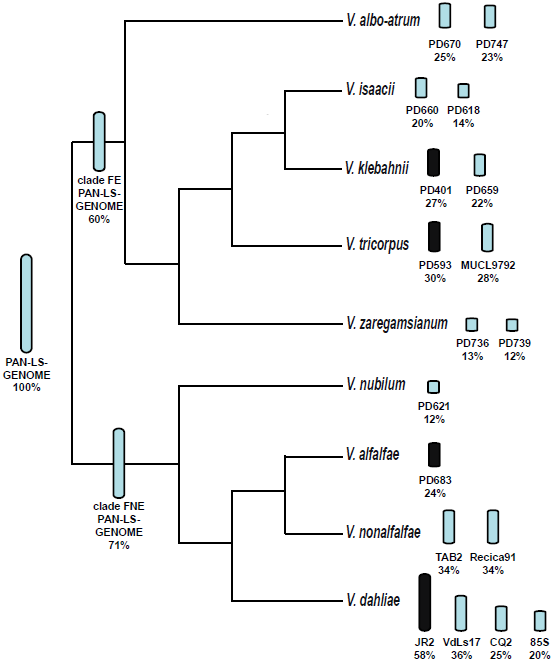
Diversity of pan-LS-genome contents across the *Verticillium* genus. A pan-LS-genome was constructed based on sequences from *Verticillium* isolates JR2, PD683, PD593 and PD401 (black bars). The bar size next to the species names in the *Verticillium* phylogenetic tree is representative for the amount of the pan-LS-genome that is present in the individual isolates. All isolates of the clade Flavexudans (FE in figure) in this study were used to calculate the percentage of the pan-LS-genome that is present in clade Flavexudans. Similarly, the portion of the Flavnonexudans (FNE in figure) in the pan-LS-genome was calculated with all isolates of the clade Flavnonexudans used in this study.

### Increased sequence conservation is not driven by negative selection

The depletion of sequence polymorphisms in LS regions may be driven by negative selection on genes with particular functions that happen to reside in these regions. Hence, we screened LS regions in the genome of *V. dahliae* strain JR2 for enrichment of particular protein domains (Pfam). In total, 13 Pfam domains with various functions are enriched in the LS region genes (Supplemental_Table_S7). However, if the negative selection on particular LS genes is responsible for observed increased sequence conservation, depletion of polymorphisms should only concern protein-coding sequences. To test this hypothesis, we compared the sequence identity of coding and intergenic regions between *V. dahliae* and *V. nonalfalfae*, which revealed that increased sequence conservation is also observed in intergenic regions (Figure 6), indicating that increased sequence conservation is likely not driven by negative selection acting on protein-coding genes.

**Figure 6:**
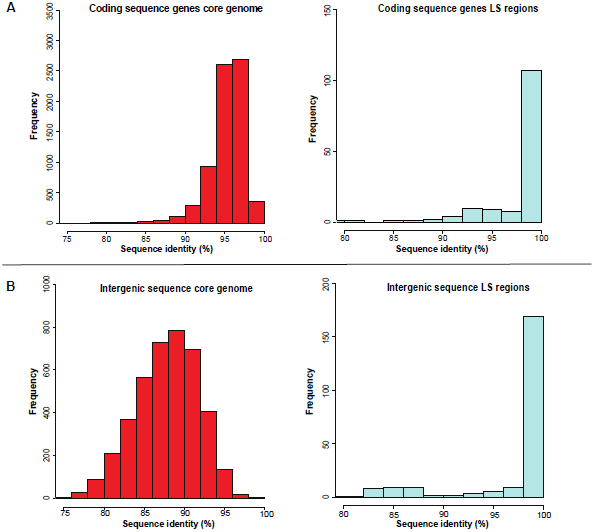
Sequence identity of *V. dahliae* JR2 core and lineage-specific (LS) regions with *V. nonalfalfae* (TAB2) for coding and intergenic sequences. (A) Coding sequence of JR2 genes were aligned to coding sequences of TAB2 genes. Matching coding sequences of genes that minimally covered 80% of each other were selected and sequence identity between their homologs was determined. (B) For the intergenic regions, windows of 5 kb were constructed for JR2 core and LS regions. The sequence identity distribution is significantly different between core and LS regions and this for both the coding sequence of genes and intergenic regions (two-sided Wilcoxon rank-sum test, *p*<0.0001).

To see how selection impacts the evolution of LS region genes, we determined the rates of non-synonymous (*Ka*) and synonymous (*Ks*) substitutions for genes that reside in LS regions versus those that reside in the core genome. In total, 49% (70 out of 142) of the LS genes could not be used for *Ka* and *Ks* determination, as we did not observe any substitutions when compared to their corresponding *V. nonalfalfae* orthologs. In contrast, within the core almost all genes (8,583 out of 8,584) display nucleotide substitutions with their respective *V. nonalfalfae* orthologs. The *Ka* was not different (two-sided Wilcoxon rank-sum test, *P*<0.05) between LS (median=0.015, *n*=74) and core (median=0.015, *n*=8583) genes (Figure 7). In contrast, the *Ks* of LS genes (median=0.12, *n*=74) was significantly lower than of core genes (median=0.16, *n*=8583). Consequently, LS genes (median=0.38, *n*=60) have significantly higher *Ka/Ks* values than core genes (median=0.09, *n*=8289), calculated for genes that have both synonymous and non-synonymous substitutions compared with their *V. nonalfalfae* orthologs. In total, 15 of the 74 tested genes displayed *Ka/Ks*>1, which is a higher proportion than the 100 of the 8,583 core genes with *Ka/Ks*>1 (Fisher’s exact test, *P*<0.05). Two LS and two core genes with *Ka/Ks*>1 were predicted to contain an N-terminal signal peptide, which is a typical characteristic of an effector protein. However, due to the limited sequence divergence in the LS regions, positive selection on the genes with *Ka/Ks*>1 was not significant based on a *Z*-test, whereas in the core genome 21 genes were found to be under positive selection (*P*<0.05). In conclusion, despite increased sequence conservation, genes in LS regions display symptoms of more diversifying selection than the core genome as *Ka/Ks* ratios were significantly higher for LS region genes.

**Figure 7:**
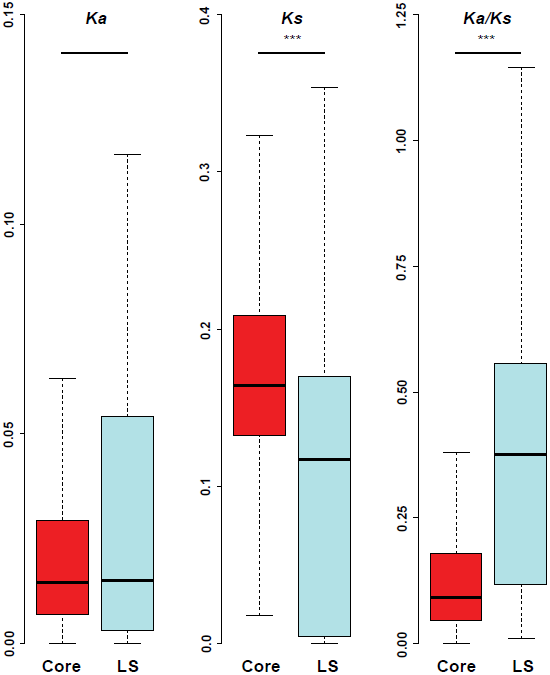
Comparison of substitutions of *V. dahliae* (JR2) and *V. nonalfalfae* (TAB2) orthologs between core and lineage-specific (LS) regions. The distribution of non-synonymous substitution rates (*Ka*), synonymous substitution rates (*Ks*) and *Ka*/*Ks* ratios are depicted for *V. dahliae* genes aligned to *V. nonalfalfae* orthologs. Outliers are not depicted. Significant differences between core and LS genes are indicated by **\*\*\*** (two-sided Wilcoxon rank-sum test, *p*<0.05).

## DISCUSSION

Genomes of many filamentous plant pathogens are thought to obey to the two-speed evolution model (Croll and McDonald 2012; Dong et al. 2015; Möller and Stukenbrock 2017). Similarly, *V. dahliae* has been suggested to evolve under a two-speed regime, as LS regions display signs of accelerated evolution as they are hot-spots for structural variation and TE activity (de Jonge et al. 2013; Faino et al. 2015; Faino et al. 2016). Additionally, we establish here that LS regions display abundant presence/absence polymorphisms between closely and even distantly related *V. dahliae* strains (Figure 1, Supplemental_Figure_S1) (de Jonge et al. 2013; Faino et al. 2016). Intriguingly, however, genomic sequences that are present within LS regions display increased sequence conservation when compared with other *Verticillium* spp. (Figure 2,3,S2; Table 1). Generally, sequences with increased identities between distinct taxa can originate from horizontal transfer, a phenomenon that has been implicated in the pathogenicity of various filamentous plant pathogens (Soanes and Richards 2014). For instance, *Pyrenophora tritici-repentis*, the causal agent of wheat tan spot, acquired a gene from the fungal wheat pathogen *Phaeosphaeria nodorum* enabling the production of the host-specific toxin ToxA that mediates pathogenicity on wheat (Friesen et al. 2006). However, the increased sequence identity of LS sequences in *V. dahliae* is likely not a consequence of horizontal transfer as homologous sequences are found in all *Verticillium* spp., and the degree of sequence conservation with these *Verticillium* spp. corresponds to their phylogenetic distance to *V. dahliae* (Table 1). Intriguingly, the increased sequence conservation is not a consequence of negative selection on coding regions, as intergenic regions display similarly increased levels of sequence conservation (Figure 6). In addition, genes residing in LS regions display higher *Ka/Ks* ratios compared to core genes, indicating the higher diversifying selection acting on these genes (Figure 7) (Sperschneider et al. 2015). In conclusion, the increased sequence conservation is likely caused by lower mutation rates in LS regions, as horizontal DNA transfer and negative selection are unlikely explanations.

Lower levels of synonymous substitutions were similarly found in the repeat-rich dispensable chromosomes of the fungal wheat pathogen *Zymoseptoria tritici* (Stukenbrock et al. 2010). However, this observation was not attributed to lower mutation rates, but rather the consequence of a lower effective population size of these dispensable chromosomes (Stukenbrock et al. 2010). Increased sequence conservation caused by lower substitution rates as observed in our study is an unprecedented for repeat-rich regions of filamentous pathogens. In contrast, previously increased substitution rates have often been associated with the two-speed genome evolution (Cuomo et al. 2007; Dong et al. 2015). For example, repeat-induced point (RIP) mutagenesis increases sequence divergence of particular effector genes of the oilseed rape pathogen *Leptosphaeria maculans* that are localized adjacent to TEs (van de Wouw et al. 2010). However, accelerated evolution through SNPs is not consistently observed for all two-speed genomes, as no significant difference in SNP frequencies between core and repeat-rich genomic regions was found for *P. infestans* (Raffaele et al. 2010).

Increased sequence conservation of LS regions seems counter-intuitive in the light of the two-speed genome model as increased variation of pathogenicity-related genes facilitates rapid evasion of host immunity. Nonetheless, an effector gene subjected to increased sequence conservation stood the test of time. The *Ave1* effector gene resides in an LS region of *V. dahliae* strain JR2 and is highly conserved as an identical copy was found in *V. alfalfae* strain VaMs102, a strain with an average nucleotide identity of 92% (de Jonge et al. 2012). Moreover, no *Ave1* allelic variation is hitherto found in the *V. dahliae* population as well as in *V. alfalfae* and *V. nonalfalfae* (de Jonge et al. 2012; Song et al. unpublished data). Conceivably, sequence conservation makes an effector gene an easy target for recognition by the host and also Ave1 is a target for immunity recognition by tomato receptor Ve1 (Fradin et al. 2009). Thus, effector catalogue diversification must be achieved through different means. Indeed, instead through SNPs, *V. dahliae* alters its effector repertoires through presence/absence polymorphisms (de Jonge et al. 2013), leading to large diversities in LS region contents between strains (Figure 5, Supplemental_Table_S6). Hence, the evasion of Ve1-mediated recognition in tomato is exclusively observed through the absence of the *Ave1* gene in *V. dahliae* strains (de Jonge et al. 2012).

Mechanisms that can explain the observed lower SNP frequency rates locally, in repeat-rich genomic regions, remain unknown. SNPs often originate from the wrong nucleotide insertion by DNA polymerase and there is no immediate reason why this should be different in LS regions. Possibly, the depletion of SNPs can be associated with a differential epigenetic organisation of LS regions, as repeat-rich regions in other filamentous pathogens are associated with densely organised chromatin, referred to as heterochromatin (Galazka and Freitag 2014). For instance, the repeat-rich conditionally dispensable chromosomes of *Z. tritici* are enriched for histone modifications associated with heterochromatin, in contrast to core chromosomes that were largely euchromatic, i.e. transcriptionally active (Schotanus et al. 2015). The link between chromatin and structural variation is under debate and controversial (Seidl et al. 2016). In general, heterochromatin is thought to suppress genomic structural alterations as recombination is repressed in heterochromatic regions of many eukaryotic genomes. However, heterochromatic regions in the filamentous pathogens *Z. tritici* are enriched for structural variations as they are enriched for duplications and deletions (Seidl et al. 2016). *V. dahliae* LS regions display similar features with enrichment of repeats, segmental duplications and presence/absence polymorphisms, hence LS regions can be anticipated to be heterochromatic (Figure 1) (de Jonge et al. 2013; Faino et al. 2016). Further research is needed to investigate whether differences in chromatin organisation may affect SNP frequencies in filamentous pathogens in general, and may explain lower rates of SNP frequencies in *V. dahliae* LS regions.

## Conclusion

The two-speed genome is an intuitive evolutionary model for filamentous pathogens, as genes important for pathogenicity benefit from frequent alternations to guarantee the continuation of symbiosis with the host. However, filamentous pathogens comprise a heterogeneous group of organisms with diverse lifestyles (Dean et al. 2012; Kamoun et al. 2015). Consequently, it is not surprising that accelerated evolution is driven by different mechanisms between species. Moreover, not all filamentous pathogens appear to adhere to the two-speed genome model (Derbyshire et al. 2017). In *V. dahliae*, acceleration evolution is merely achieved through presence/absence polymorphisms, as nucleotide sequences are highly conserved in LS regions. The dependency of host adaptation on presence/absence polymorphisms may lead to a more rapid immunity evasion than sequence alterations through SNPs (Daverdin et al. 2012). Thus, the quick fashion of host immunity evasion through the deletion of effector genes can be evolutionary advantageous over allelic diversification, especially for pathogens with a small effective population size.

## METHODS

### Genome sequencing and assembly of *Verticillium* isolates

In total, we used 18 *Verticillium* genomes in this study (Supplemental_Table_S2). Genomes of *V. albo-atrum* PD747, *V. alfalfae* PD683, *V. dahliae* JR2 and VdLs17, *V. isaacii* PD618, *V. klebahnii* PD401, *V. nubilum* PD621, *V. tricorpus* PD593 and MUCL9792, *V. zaregamsianum* PD739 were previously sequenced and assembled (Klosterman et al. 2011; Faino et al. 2015; Seidl et al. 2015; Shi-Kunne et al. in preparation). Furthermore, sequence reads of the two *V. nonalfalfae* isolates (TAB2 and Rec) were publically available (Bioproject PRJNA283258) (Jelen et al. 2016). *Verticillium* strains CQ2, 85S, PD670, PD660, PD659 and PD736 were sequenced in this study. To this end, we isolated genomic DNA from conidia and mycelium fragments that were harvested from cultures that were grown in liquid potato dextrose agar according to the protocol described by Seidl *et al.* (2015). We sequenced *V. dahliae* strains CQ2 and 85S by single molecule real time (SMRT) sequencing. The PacBio libraries for sequencing on the PacBio RSII platform (Pacific Biosciences of California, CA, USA) were constructed as described previously by Faino et al. (2015). Briefly, DNA was mechanically sheared and size selected using the BluePippin preparation system (Sage Science, Beverly, MA, USA) to produce ~20 kb size libraries. The sheared DNA and final library were characterized for size distribution using an Agilent Bioanalyzer 2100 (Agilent Technology, Inc., Santa Clara, CA, USA). The PacBio libraries were sequenced on four SMRT cells per *V. dahliae* isolate using the PacBio RS II instrument at the Beijing Genome Institute (BGI, Hong Kong) for CQ2 and at KeyGene N.V. (Wageningen, the Netherlands) for 85S, respectively. Sequencing was performed using the P6-C4 polymerase-Chemistry combination and a >4 h movie time and stage start. Filtered sub-reads for CQ2 and 85S, were assembled using the HGAP v3 protocol (Supplemental_Table_S1) (Chin et al. 2013).

For PD670, PD660, PD659 and PD736, two libraries (500 bp and 5 Kb insert size) were prepared and sequenced using the Illumina High-throughput sequencing platform (KeyGene N.V., Wageningen, The Netherlands). In total, ~18 million paired-end reads (150 bp read length; 500 bp insert size library) and ~16 million mate-paired read (150 bp read length; 5 kb insert size library) were produced per strain. We assembled the genomes using the A5 pipeline (Tritt et al. 2012), and we subsequently filled the remaining sequence gaps using SOAPdenovo2 (Luo et al. 2012). After obtaining the final assemblies, we used QUAST (Gurevich et al. 2013) to calculate genome statistics. Gene annotation for *V. dahliae* strain JR2 and other *Verticillium* spp. were obtained from Faino et al. (2015) and Shi-Kunne et al. (in preparation). Genes for *V. isaacii* strain PD660 were annotated with the Maker2 pipeline according to Shi-Kunne et al. (in preparation) (Holt and Yandell 2011).

### Comparative genome analysis

The alignments of *Verticillium* sequences to a reference genome were performed with nucmer, which is part of the mummer package (v3.1) (Kurtz et al. 2004). Here, we used a repeat-masked genome as a reference in order to prevent assigning high sequence identities to repetitive elements. Repetitive elements were identified using RepeatModeler (v1.0.8) based on known repetitive elements and on *de novo* repeat identification, and genomes were subsequently masked using RepeatMasker (v4.0.6; sensitive mode) (Smit et al. 2015).

Linear plots showing alignments within and closely adjacent JR2 LS regions were plotted with the R package genoPlotR (Guy et al. 2011) (Figure 2, Supplemental_Figure_S3). The *Verticillium* phylogenetic tree adjacent to the genoPlotR plots was previously generated using 5,228 single-copy orthologs that are conserved among all of the genomes (Shi-Kunne et al. in preparation). The phylogenetic tree of *V. dahliae* strains was constructed using REALPHY (Bertels et al. 2014) (Supplemental_Figure_S1).

Alignments > 7.5 kb in length were depicted along the reference genome with the R package Rcircos (Figure 3,4) (Zhang et al. 2013). LS sequences were defined by alignment of different strains to a reference using nucmer (v3.1) (Kurtz et al. 2004) and regions were determined using BEDTools v2.25.0 (Quinlan and Hall 2010).

Lineage-specific regions of *V. dahliae* and *V. tricorpus* were arbitrarily delimited based on the abundance of LS sequences and increased sequence conservation (Supplemental_Table_S8). The pairwise identity of the genome-wide and LS regions between *V. dahliae/V. tricorpus* and other haploid *Verticillium* spp. was calculated using nucmer (mum), with dividing the respective query sequences into non-overlapping windows of 500 bp (Table 1). Sequence identities of the coding regions of genes and intergenic regions were retrieved by BLAST (v2.2.31+) searches between strains *V. dahliae* JR2 and *V. nonalfalfae* TAB2 (Figure 6) (Altschul et al. 1990). Hits with a minimal coverage of 80% with each other were selected. Intergenic regions of JR2 were fractioned in 5 kb windows with BEDTools v2.25.0 and similarly blasted to the genome of TAB2 (Figure 6) (Quinlan and Hall 2010). Hits with a maximal bit-score and minimal alignment of 500 bp to a window were selected. To compare the rate of synonymous and non-synonymous substitutions between the core and LS regions, *Ka* and *Ks* were of orthologs of JR2 and TAB2 were determined using the Nei and Gojobori method (Nei and Gojoborit 1986) in PAML (v4.8) (Yang 2007). Significance of positive selection was tested using a Z-test (Stukenbrock and Dutheil 2012). Z-values >1.65 were considered significant with *P*<0.05. Secreted proteins were predicted by SignalP4 (Petersen et al. 2011).

Pfam function domains of JR2 proteomes were predicted using InterProScan (Jones et al. 2014). Subsequently, Pfam enrichment of genes residing in LS regions was carried out using hypergeometric tests, and significance values were corrected using the Benjamini-Hochberg false discovery method (Benjamini and Hochberg 1995).

The pan-LS-genome was constructed based on following *Verticillium* isolates: JR2 (*V. dahliae*), PD683 (*V. alfalfae*), PD593 (*V. tricorpus*) and PD401 (*V. klebahnii*). Genome regions of these for species with increased sequence conservation were combined (Supplemental_Table_S8). Repeat masked regions were removed from the pan-LS-genome using BEDTools v2.25.0 (Quinlan and Hall 2010). Additionally, regions in duplicate (≥90% identity, ≥100bp) in the pan-LS-genome were determined using nucmer (v3.1) (Kurtz et al. 2004) and subsequently removed with using BEDTools v2.25.0 (Quinlan and Hall 2010). As result, a pan-LS-genome was constructed without regions in duplicate. The fractions of pan-LS-genome that were present in every individual *Verticillium* strain was determined using nucmer (v3.1) (Kurtz et al. 2004). The clade pan-LS-genomes were constructed by combining all the pan-LS-genome regions that are present in the *Verticillium* clade isolates, which was then also removed from duplicate regions.

## ACKNOWLEDGEMENTS

The authors would like to thank the Marie Curie Actions program of the European Commission that financially supports the research of J.R.L.D. Work in the laboratories of B.P.H.J.T. and M.F.S is supported by the Research Council Earth and Life Sciences (ALW) of the Netherlands Organization of Scientific Research (NWO). H. M. was supported by French Ministry of Higher Education and Research: CIFRE 2013/1431.

## DATA ACCESS

The Whole Genome Shotgun projects have been deposited at DDBJ/ENA/GenBank as accessions PRLI00000000 and PRLJ00000000 for *V. dahliae* strains CQ2 and 85S, respectively.

